# Local adaptation can cause both peaks and troughs in nucleotide diversity within populations

**DOI:** 10.1101/2020.06.03.132662

**Authors:** Russ J. Jasper, Sam Yeaman

## Abstract

Local adaptation is expected to cause high F_ST_ at sites linked to a causal locus, however this pattern can also be driven by background or positive selection. Within-population nucleotide diversity could provide a means to differentiate these scenarios, as both background and positive selection deplete diversity, whereas some theoretical studies have shown that local adaptation increases it. However, it is unclear whether such theoretical predictions generalize to more complicated models. Here, we explore how local adaptation shapes genome-wide patterns in nucleotide diversity and F_ST_, extending previous work to study the effect of variable degrees of polygenicity and genotypic redundancy in an adaptive trait, and different levels of population structure. We show that local adaptation produces two very different patterns depending on the relative strengths of migration and selection, either markedly decreasing or increasing within-population diversity at linked sites at equilibrium. When migration is low, regions of depleted diversity can extend large distances from the causal locus, with substantially more diversity eroded than expected with background selection. With higher migration, peaks occur over much smaller genomic distances but with much larger magnitude changes in diversity. In spatially extended clinal environments both patterns can be found within a single species, with increases in diversity at the center of the range and decreases towards the periphery. Our results demonstrate that there is no universal diagnostic signature of local adaptation based on nucleotide diversity, however, given that neither background nor positive selection inflate diversity, when peaks are found they strongly suggest local adaptation.

## INTRODUCTION

Understanding how evolutionary processes shape genetic variation is crucial for interpreting patterns across the genome. Early genome scan studies identified outlier peaks in F_ST_ as putative indicators of locally adapted loci (reviewed in Via 2009; Nosil et al. 2009), because strong selection is expected to drive high differentiation in allele frequency at selected loci (Lewontin and Krakauer 1973) and linked neutral sites (Charlesworth et al. 1997, Feder and Nosil 2010). Subsequent reinterpretation of these patterns, however, suggested that outlier peaks in F_ST_ could also be generated by reductions in within-population nucleotide diversity driven by “linked selection” (Noor and Bennett 2009; Cruickshank and Hahn 2014), either as a result of recent selective sweeps (Maynard Smith and Haigh 1974; Kaplan et al. 1989; Stephan et al. 1992; Braverman et al. 1995; Gillespie 1997, 2000, 2001) or background selection (Charlesworth et al. 1993; Charlesworth 1994; Hudson and Kaplan 1995; Gillespie 1997). Furthermore, some studies have demonstrated that genome-wide patterns in F_ST_ tend to be inversely correlated with both recombination rate and nucleotide diversity, suggesting that the search for the causal loci driving local adaptation may be obfuscated by the recombination and/or diversity landscapes across the genome (Burri et al. 2015; Vijay et al. 2017; Irwin et al. 2018).

In light of these studies, it is now well-recognized that both hard sweeps and background selection have the potential to reduce genetic variation at linked sites (Booker et al. 2019), and they are now commonly invoked to explain signatures where a relative local reduction in nucleotide diversity is found in some areas of the genome along with elevated F_ST_. Complicating the picture somewhat, recent theoretical work has suggested that background selection likely only very minimally affects F_ST_ and that any detected peaks are therefore unlikely to be generated solely as a function of reduced diversity due to background selection (Matthey-Doret and Whitlock 2019). Still other studies have shown that uniform positive selection can generate high among-population variation as a consequence of incomplete sweeps or recombination during a sweep, which may be sufficient to explain observed genome-wide patterns of F_ST_ in many cases (Bierne 2010; Booker et al. 2019).

Regardless of what drives genome-wide patterns of F_ST_, it remains that local adaptation *does* often occur (Hedrick et al. 1976; Linhart and Grant 1996; Hereford 2009), and as such, some signatures of elevated F_ST_ may also be expected in species with local adaptation. Given that within-population nucleotide diversity is now commonly used to infer the possible activity of background or positive selection at linked sites, it is also important to understand how local adaptation affects nucleotide diversity, especially as it pertains to interpreting genome scan results. Note: hereafter, we will use the term “nucleotide diversity” to refer to the within-population component, unless specifically stated otherwise.

For an unlinked neutral locus in an island model, expected nucleotide diversity is 4*N*_*e*_μ when migration rate is 0 and increases with *m* (and scales with 1 – F_ST_) towards 4*dN*_*e*_μ at panmixia (where *N*_*e*_ is effective population size, μ is the per-locus mutation rate, *m* is migration rate, and, *d* is the number of demes). For a neutral locus linked to a divergently-selected site, it is well-known that the effective migration rate the neutral locus experiences decreases with increased linkage (Petry 1983, Bengtsson 1985; Barton and Bengtsson 1986). Thus, at low migration rates, nucleotide diversity near a selected site should also be depleted due to the lower effective population size, which can also be shown by rearranging results from Petry (1983, Eq. 14) to yield a prediction for expected heterozygosity (H_S_) of

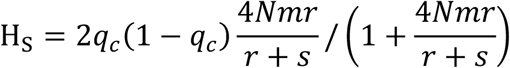

where *q*_*c*_ is the allele frequency at the neutral locus in the continental population, *N* is the population size, *m* is the migration rate, *r* is the recombination rate, *s* is the selection coefficient, and given

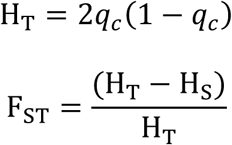

This rearranged equation for H_S_ (Petry 1983) predicts a monotonic decrease in nucleotide diversity with decreasing recombination or migration. Nordborg (1997) likened the effect of migration-selection balance in this range of parameter space to that of background selection, which also decreases diversity at linked sites (also see Aeschbacher and Bürger 2014; Fig. 8).

It is less clear, however, what happens to nucleotide diversity as migration rate increases, and the above models make assumptions that limit their applicability to high migration. It is well accepted that strong balancing selection increases total nucleotide diversity at linked sites (Hudson and Kaplan 1988; Barton and Navarro 2002; Charlesworth 2006), and divergent selection with high migration would therefore have similar effects (Nordborg and Innan 2003). It is unclear, however, how such diversity would be partitioned within vs. among populations and how this would change with migration-selection balance. The set of individuals with a given divergently selected haplotype can be considered analogous to a population, and we would expect reduced diversity around a selected site within this set, and increased diversity among sets. However, while selection acts to increase the assortment of locally adapted haplotypes to the population where they are favoured, migration mixes haplotypes and inflates diversity, and as such, increased within-population diversity might be expected. Indeed, Charlesworth et al. (1997) showed an increase in within-population nucleotide diversity near the locally adapted locus for a model that also included background selection (their Figure 7A), and for an analytical model without background selection (their Table 1). More recently, Sakamoto and Innan (2019) analyzed a two-population model and found a small peak in within-population nucleotide diversity around the locally adapted locus, but focused more of their analysis on the decrease in diversity around the selected site that occurs during the initial establishment of a locally adapted polymorphism. As no study seems to have directly and broadly focused on the effect of migration-selection balance on within-population nucleotide diversity at linked sites, further consideration of this question is necessary.

**Table 1:**
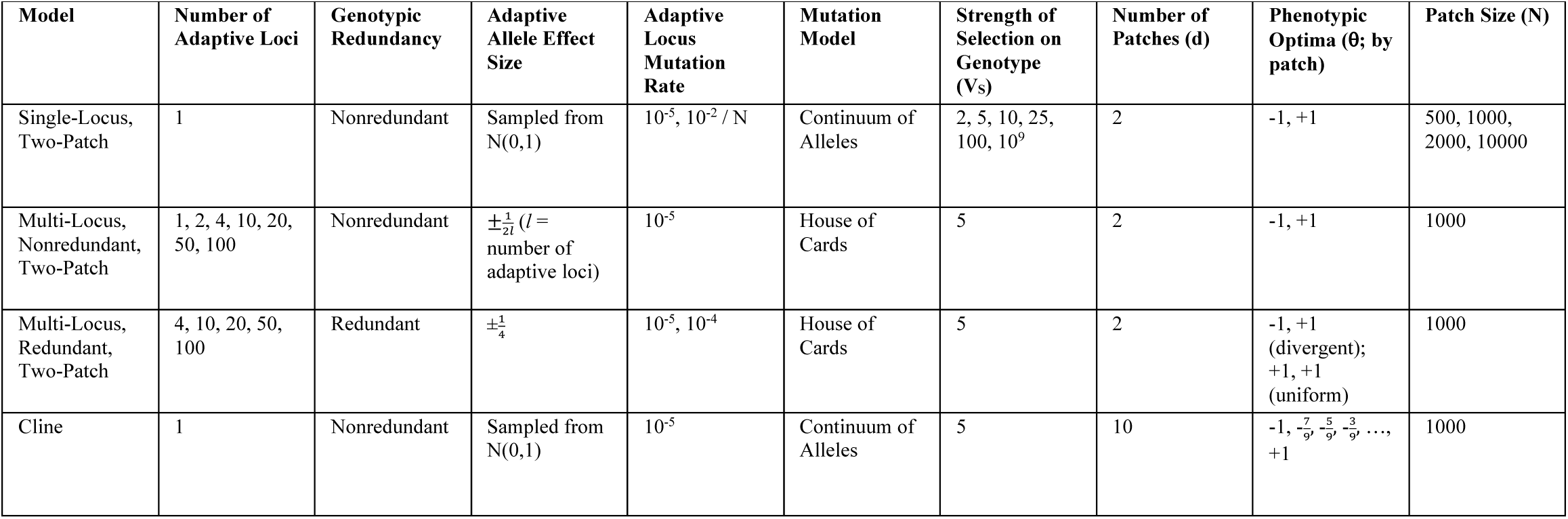
Model parameters and associated values.

In the present study, we describe how local adaptation over heterogenous landscapes shapes patterns in nucleotide diversity and F_ST_ at the neutral regions flanking a selected locus over a wide range of migration-selection parameter space. We begin by using individual-based simulations of two-patch models with a single selected locus, then explore models with polygenic adaptive traits and varying degrees of genotypic redundancy, and lastly, investigate more complex patterns of population structure by exploring a ten-patch stepping stone cline. We show that both troughs and peaks in nucleotide diversity may be expected around a locally-adapted locus, depending on the parameters involved.

## MATERIALS AND METHODS

To understand how local adaptation over heterogenous landscapes shapes patterns in nucleotide diversity we performed simulations using two different landscape models: a two-patch model (with and without genotypic redundancy) and a linear ten-patch cline model (Table 1). We performed our simulations with the stochastic, forward-time, individual-based simulation program Nemo, version 2.3.46 (Guillaume and Rougemont 2006). Our simulations followed the Wright-Fisher model with the addition of selection, migration, and mutation. We modelled a single trait under Gaussian stabilizing selection, where the fitness of an individual (*W*) was defined as

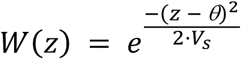

Where *z* was an individual’s phenotypic value, an additive function of the allele effect size at each locus (i.e. no epistasis or dominance effects on phenotype); θ was the optimal phenotypic value of the local patch; and V_S_ was the strength of selection on the genotype as described by the variance around the fitness function. Unless otherwise stated, a V_S_ of 5 was used. After calculating the base fitness value of each individual within a patch, an individual’s fitness was scaled against the mean fitness of the local patch.

Individuals had a single diploid chromosome where each divergently selected locus was symmetrically flanked by 74 neutral loci positioned at distances from 10^−3^ to 10 cM away on a log_10_ scale (Fig. S1 & S2). In the case of multiple adaptive loci on a chromosome, each adaptive locus and its 74 flanking neutral loci was separated from the next closest adaptive locus and associated neutral loci by 50 cM, such that one complement of 75 loci was unlinked from any other complement of 75 loci (Fig. S2). Neutral loci were diallelic and mutation occurred at a rate of 10^−5^ per locus per generation. Similarly, unless otherwise stated, mutation occurred at the selected loci at a rate of 10^−5^.

Forward migration rates were varied between 10^−5^ to 10^−½^ with four equal increments per order of magnitude, in addition to a migration rate of zero. Unless otherwise stated, each patch was comprised of 1,000 individuals. Each simulation replicate was run for a total of 25·N·d generations, where N was the population size by patch and d was the number of patches in the metapopulation. The within- and between-population nucleotide diversity, total metapopulation nucleotide diversity and F_ST_ were then calculated at each locus after 25·N·d generations, except in the case of the single-locus, two-patch model, where they were iteratively calculated every ½·N·d generations.

### Single-Locus, Two-Patch Model

In the single-locus, two-patch model, the adaptive locus was multiallelic and mutation occurred whereby a new allele effect size was drawn from a normal distribution (μ = 0, σ^2^ = 1) and added to the former allele effect size (i.e. continuum of alleles). Two different mutation rates were explored, an unscaled rate independent of population size (10^−5^) and a scaled rate standardized by population size (10^−2^ / N) (i.e. populations had the same number of mutational events per generation regardless of population size).

### Multiple Adaptive Loci, Two-Patch Model

In models with polygenic adaptive traits, adaptive loci were diallelic and mutation occurred whereby the original allele would be replaced by the opposite allele (i.e. house of cards). In the genotypically nonredundant model, the allele effect sizes were scaled relative to the number of adaptive loci such that an individual needed to be homozygous for the optimal allele at every locus in order to achieve the optimal phenotype in a given patch. By contrast, in the genotypically redundant model, the allele effect sizes were set to ± 0.25 regardless of the number of loci, such that an individual could reach the phenotypic optimum (± 1 divergent selection; + 1 uniform selection) by being homozygous for the optimal alleles at any two loci.

### Single-Locus, Cline Model

Our cline model consisted of ten patches in a linear conformation and followed the stepping stone migration model (i.e. dispersal was only possible between directly adjacent patches). The adaptive locus was multiallelic and mutation occurred as in the single-locus, two-patch model, where a new allele effect size was drawn from a normal distribution (μ = 0, σ^2^ = 1) and added to the former allele effect size (i.e. continuum of alleles).

### Study Metrics

Nucleotide diversity (π) was calculated as

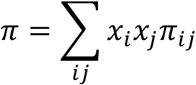

Where *x*_*i*_ and *x*_*j*_ were the respective frequencies of the *i*^th^ and *j*^th^ sequences in a population and *π*_*ij*_ was the number of nucleotide differences between the *i*^th^ and *j*^th^ sequences (Nei and Li 1979). As well, we report the F_ST_ per locus returned by Nemo, which was calculated using Weir and Cockerham (1984). We regressed the nucleotide diversity or F_ST_ at a neutral locus on its log_10_-transformed distance from an adaptive locus (cM) to quantify the relationship between diversity or F_ST_ and distance from an adaptive locus. As an additional way to explore the effect of selection and linkage on diversity, we compared the levels of nucleotide diversity or F_ST_ at the neutral loci 0.001 cM away from the adaptive locus to those 10 cM away.

In order to study the effect of selection across the chromosome, we calculated the mean level of nucleotide diversity that persisted under strong selection (V_S_ of 5) at all neutral loci between 9 to 10 cM away from the adaptive locus, and compared this to the genome-wide background levels of diversity across the chromosome under neutral evolution (V_S_ of 10^9^). For this specific analysis we added additional neutral loci to the ends of the chromosome, such that there were 100 loci positioned from 9 to 10 cM away from the adaptive locus, in even steps, at either end of the chromosome. The additional neutral loci were added in order to reduce the noise in our result and did not affect the overarching patterns seen.

We identified peaks in nucleotide diversity as those with a significant slope of diversity by log_10_-transformed distance according to a t-test and at least 25% of neutral loci with diversity levels in excess of 1.1 times the genome-wide background level. To quantify the width of a peak in diversity, we regressed the nucleotide diversity of all neutral loci with levels in excess of the genome-wide background on their distance from the adapted locus; the x-intercept of the regression was taken as one half of the width of the peak.

Finally, we estimated pairwise linkage disequilibrium between the adapted locus and each neutral locus across the metapopulation as a whole. Pearson’s *r*^2^ was taken as an estimate of linkage disequilibrium.

### Comparison with Previous Analytic Predictions

We compared our simulation results for the within-population nucleotide diversity at a tightly linked locus (10^−3^ cM) to the analytical predictions for the expected within-population heterozygosity from Sakamoto and Innan (2019) equation 25. Where we model Gaussian fitness acting on the phenotype (to facilitate comparison among single- and multi-locus traits), Sakamoto and Innan (2019) used selection coefficients acting on an individual locus in each patch (*s*_*i*_). To compare models, we set each of the *s* terms in eq. 25 to match the reduction in fitness for a locally optimal individual moving to the non-optimal patch (and ignored the effect of dominance), which requires asymmetrical coefficients (*s*_1_ ≠ |*s*_2_|). We also report the comparison between models under symmetrical selection coefficients.

## RESULTS

### Single-Locus, Two-Patch Model

#### Migration-Selection Balance

To explore the effect of selection on nucleotide diversity at linked sites, we calculated the slope of the regression of mean nucleotide diversity on the distance from the selected locus, which we will refer to as the diversity-distance-slope (dd-slope). When this slope was positive, nucleotide diversity tended to be substantially depressed near the selected site (as occurs with a selective sweep or background selection), whereas a peak in diversity around the selected site was present when the slope was negative. In the single-locus, two-patch model we observed a non-monotonic relationship between migration rate and the dd-slope, with slopes of zero when migration rate was zero, positive slopes at very low migration rates, and negative slopes at intermediate-high migration rates, with a transition between these opposite patterns at low-intermediate migration rates (Fig. 1a). These patterns can also be seen by contrasting the nucleotide diversity at the neutral loci nearest the adapted locus compared to those furthest away (Fig 1b). When the dd-slope was positive we found that the diversity was substantially depressed across the entire length of the chromosome (10 cM on either side of the selected locus) relative to genome-wide background levels (Fig. S3a), whereas when the dd-slope was negative we observed a much more restricted effect across the chromosome, with peaks in diversity of 0.14 – 1.18 cM in width beyond background levels (Fig. S3b).

**Figure 1:**
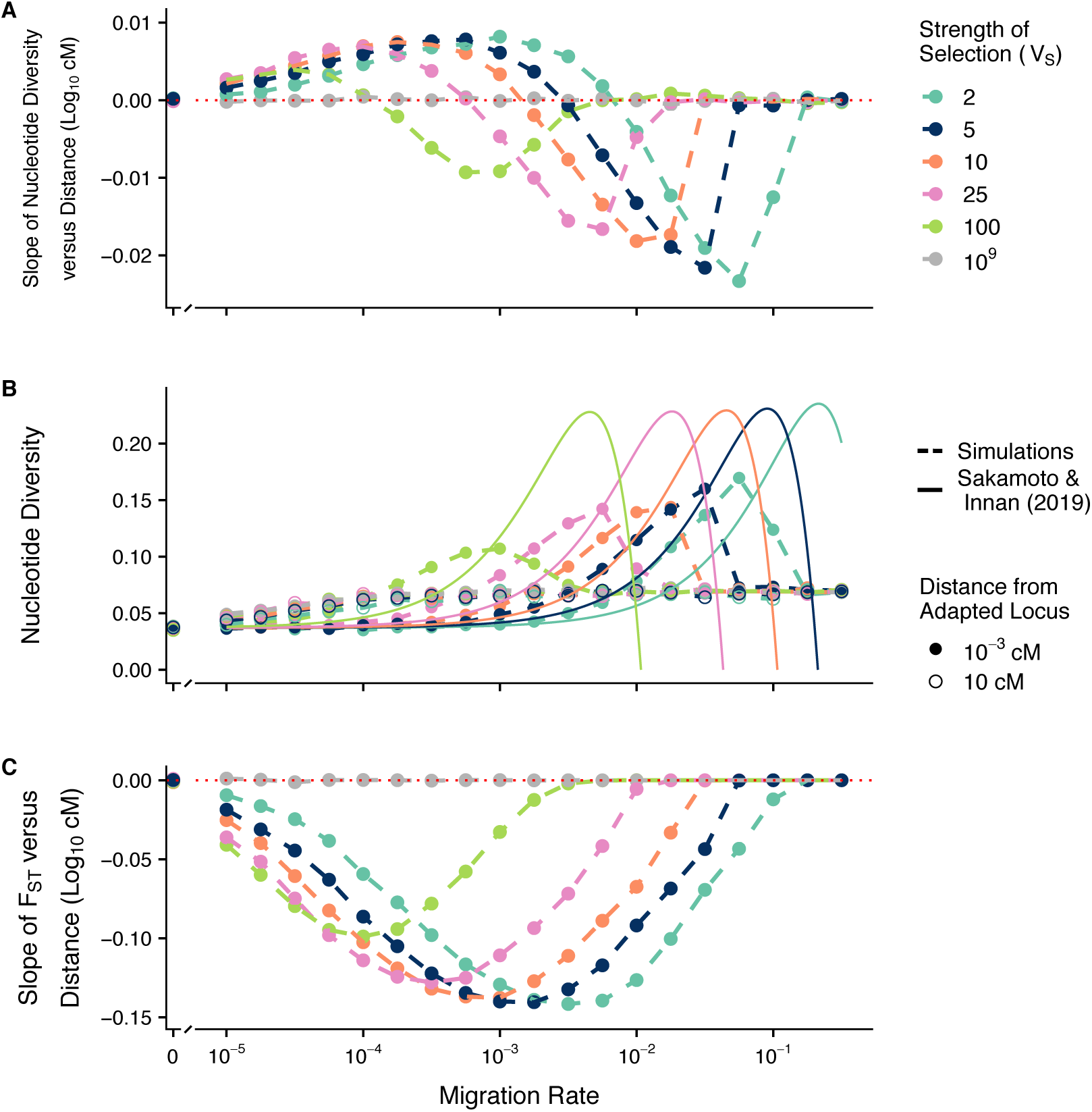
Effect of migration-selection balance on diversity at neutral sites linked to a single divergently selected locus in two-patch model. The slope of within-population nucleotide diversity versus distance (log_10_ cM) (A), the nucleotide diversity at the loci closest (10^−3^ cM) and furthest (10 cM) from the locally adapted locus (B), and the slope of F_ST_ versus distance (log_10_ cM) (C) are shown against the log_10_ migration rate after 50,000 generations. Each patch was comprised of N = 1,000 individuals and the per-locus mutation rate = 10^−5^. Simulation results are shown in dashed lines, analytical predictions from Sakamoto and Innan (2019) equation 25 using asymmetrical selection coefficients are shown in solid lines.

In cases where migration was too high to permit the maintenance of local adaptation, the dd-slope tended to return to 0 (Fig. 1a) and the per locus nucleotide diversity at tightly linked neutral loci declined to approximate the drift expectation, as indicated by the case with Vs = 10^9^ (grey line, Fig. 1b). Increasing the strength of selection shifted the above described patterns so that the peaks and transitions occurred at higher rates of migration, and also increased the maximum magnitudes of both the peak positive and peak negative dd-slopes (Fig. 1a-b).

Additionally, we used the same approach to examine patterns in total metapopulation nucleotide diversity, between-population nucleotide diversity (*d*_*xy*_) and F_ST_. As might be expected from previous theoretical work (Charlesworth et al. 1997; Sakamoto and Innan 2019), the between-population dd-slope (Fig. S4) and the slope of F_ST_ by distance (Fig. 1c) was always negative and reached a maximum magnitude at intermediate migration rates, as these conditions maximized the difference between the between-population diversity or F_ST_ at the selected locus and the same metric at unlinked neutral loci.

We found high qualitative concordance between our results for the within-population nucleotide diversity at a tightly linked locus (10^−3^ cM) and Sakamoto and Innan’s (2019) analytical prediction for the expected heterozygosity with both asymmetrical (Fig. 1b) and symmetrical nonzero selection coefficients (Fig. S5). In our simulation results, however, we did find a strong effect of the strength of selection on the maximum magnitude of the peaks in diversity observed over high migration rates. Conversely, the magnitude of the peaks in expected heterozygosity predicted from Sakamoto and Innan (2019) appear relatively insensitive to the strength of selection.

#### Effects of Population Size

We investigated how altering the population size of each patch affected the patterns in nucleotide diversity and F_ST_ described above. Broadly speaking, increasing the population size increased the value of both the dd-slopes and the slopes of F_ST_ by genetic distance over all migration rates below the critical rate (Fig. 2a,c). Over intermediate-high migration rates, increasing the patch size to 10,000 eliminated the negative dd-slope trend (Fig. 2a) and any peaks in diversity beyond the neutral expectation (Fig. 2b) seen with smaller population sizes. Across these intermediate-high migration rates, the linkage disequilibrium between neutral loci and the locally adapted locus decayed much more rapidly with larger population sizes (Fig. S6). Finally, there was little noticeable effect of scaling the mutation rate by population size (Fig. 2a,c).

**Figure 2:**
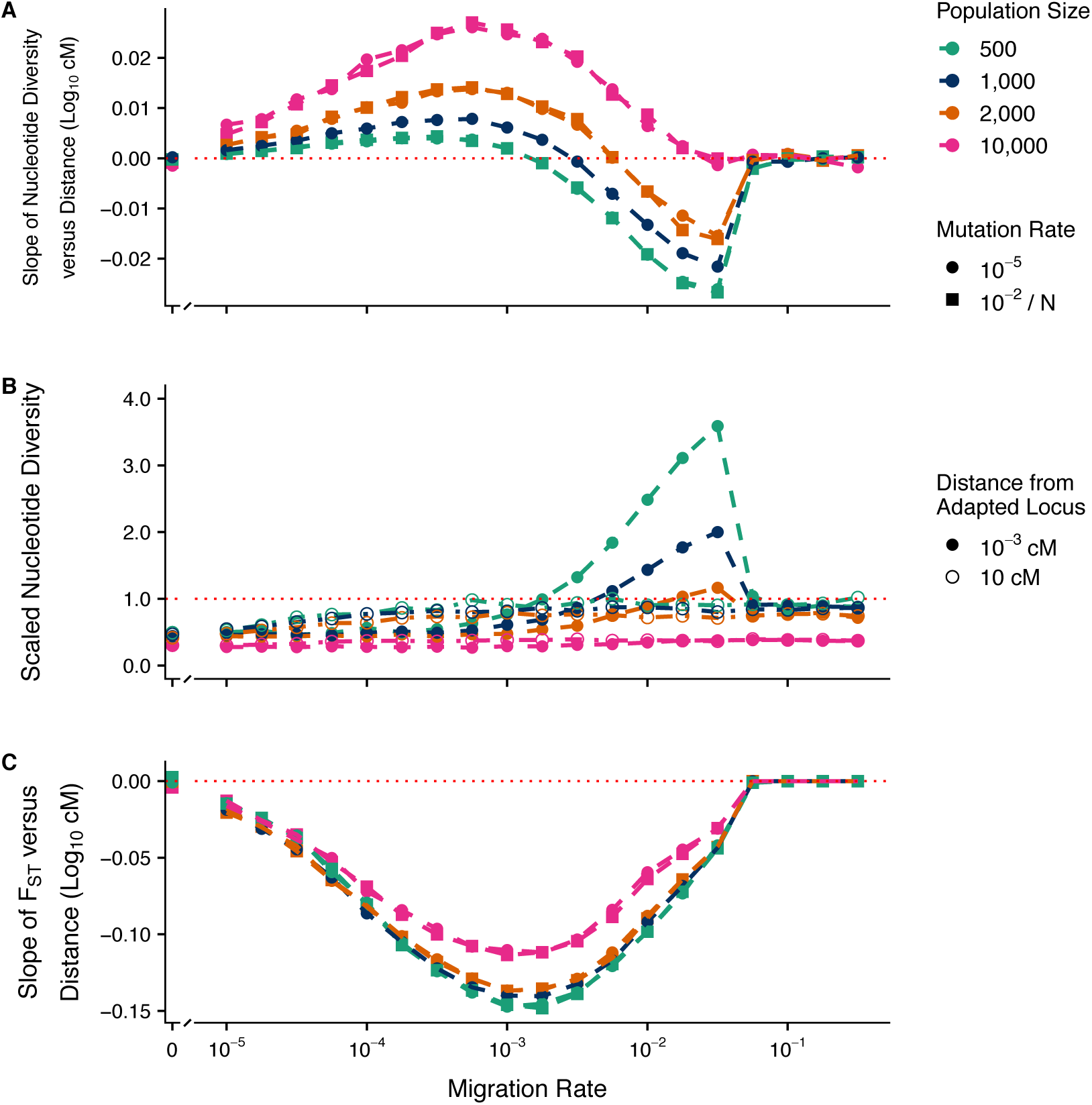
Effect of migration-selection balance and population size on diversity at neutral sites linked to a single divergently selected locus. The mutation rate at the selected locus was 10^−5^ (unscaled) or 10^−2^ / N (scaled), the mutation rate at neutral loci was 10^−5^ per locus, and V_S_ = 5. Panels A and C are as described in Fig. 1; panel B shows the nucleotide diversity scaled by the neutral expectation at *m* = 0.5 (4·d·N·μ).

#### Patterns through Time

To explore how patterns in nucleotide diversity and F_ST_ might change with time, we examined the slopes at 1000-generation intervals. Both the dd-slopes and the slopes of F_ST_ by distance consistently decreased with time until reaching their respective equilibrium values (Fig. S7a,c). Additionally, the per locus diversity steadily decreased with time until equilibrating, where the loci closest to the locally adapted locus reached equilibrium earlier and the loci furthest away later (Fig. S7b).

### Effects of Multiple Adaptive Loci and Genotypic Redundancy

When there was no genotypic redundancy (i.e. when mutations at all loci were needed to yield a locally optimal phenotype), an increase in the number of adaptive loci corresponded to a decrease in the net effect of selection on each individual locus, as each locus had a smaller allele effect size. Thus, the effect of increasing the number of loci (Fig. 3) closely resembled the effect of reducing the strength of selection observed in the single-locus model (Fig. 1). In contrast, when there was genotypic redundancy in the trait (i.e. more loci than the number of mutations needed to reach the local optimum) and each adaptive locus had the same effect size regardless of the total number of loci involved, increasing the number of loci did not shift the patterns of dd-slope with migration through the parameter space (Fig. 4).

**Figure 3:**
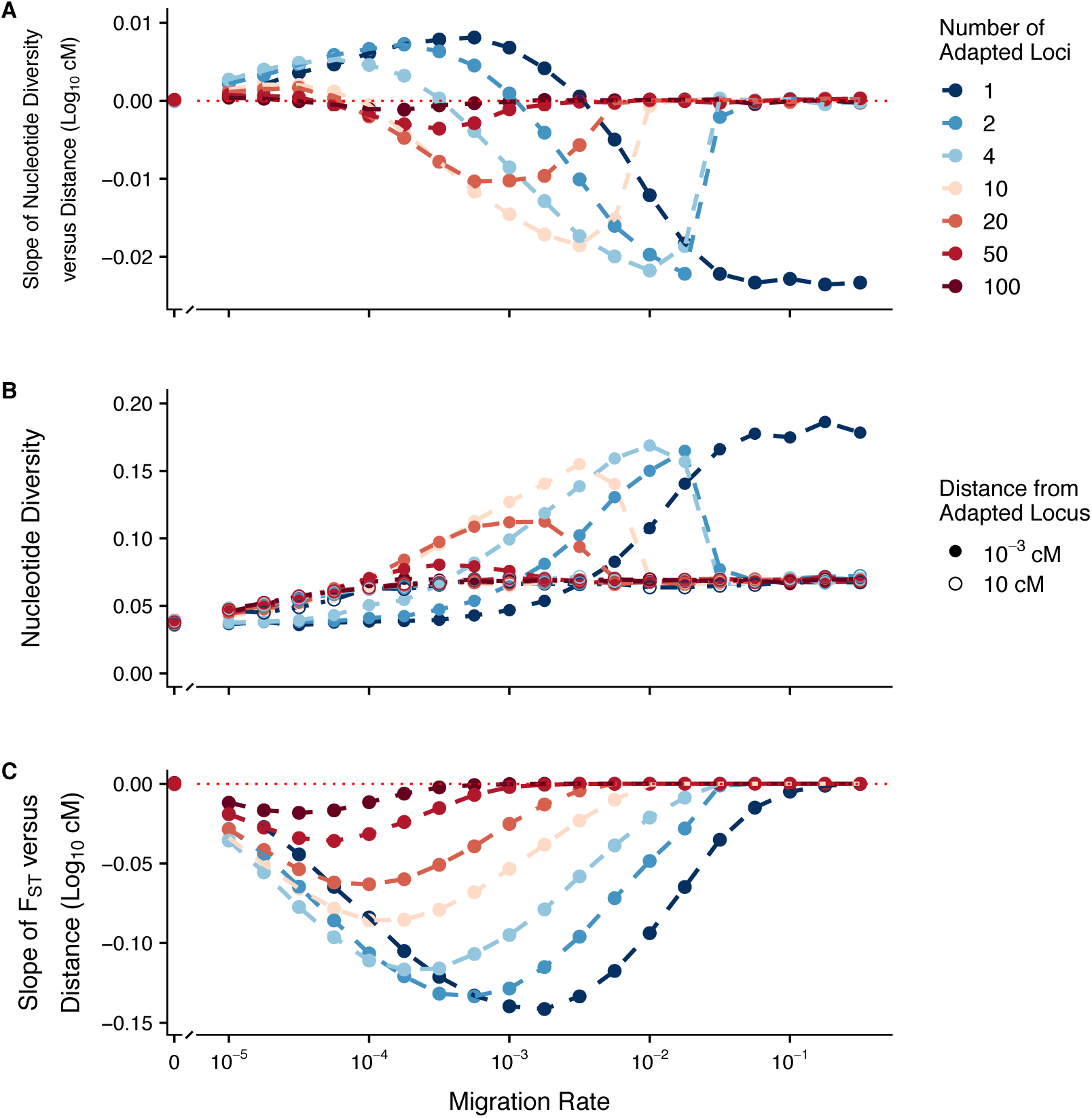
Effect of migration-selection balance on diversity at linked neutral sites for a quantitative trait with different numbers of loci and no genotypic redundancy. Allele effect sizes were scaled by the number of adaptive loci, such that an individual could only reach the optimum in a given patch by being homozygous for the optimal allele at each locus. Each patch was comprised of N = 1,000 individuals, mutation rate 10^−5^ per locus, and V_S_ = 5. Panels are as described in Fig. 1.

**Figure 4:**
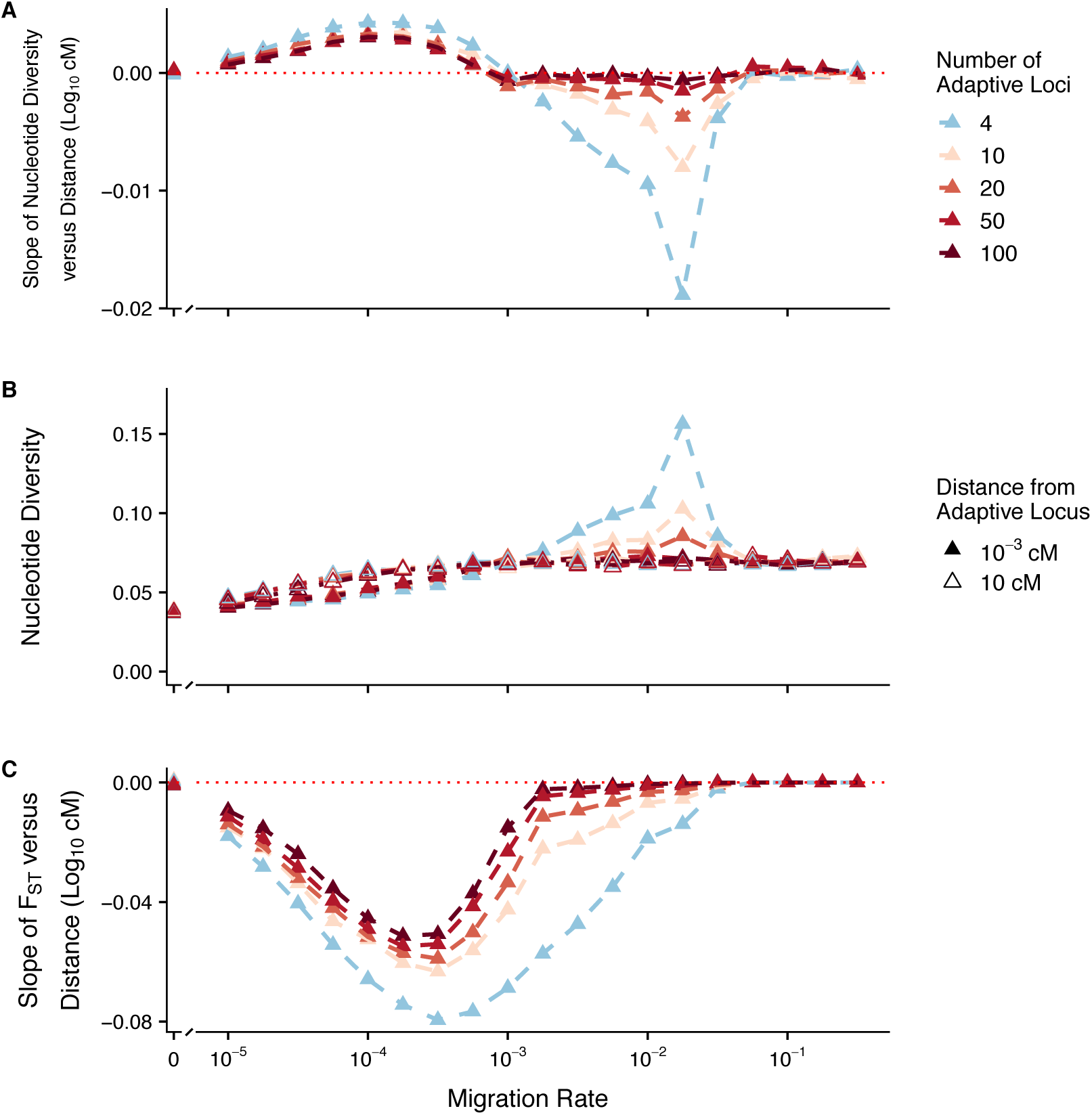
Effect of migration-selection balance on diversity at linked neutral sites for a quantitative trait with different numbers of loci and variable levels of genotypic redundancy. Allele effect sizes were ± 0.25, such that an individual could reach the optimum in a patch (± 1) by being homozygous for the optimal allele at 2 loci. Each patch was comprised of N = 1,000 individuals, mutation rate = 10^−5^ per locus, and V_S_ = 5. Panels are as described in Fig. 1.

Across low migration rates where we found reduced diversity around the focal site, we observed an interaction between the number of adaptive loci contributing to a trait and whether or not there was genotypic redundancy (Fig. 3 & 4). In the case with no redundancy, the relationship between dd-slope and migration attenuated with an increasing number of loci, with the transition point between positive and negative slopes occurring at progressively lower migration rates with increasing number of loci (Fig. 3). By contrast, in the case with redundancy, there was an attenuation in the increase in diversity found at high migration rates with an increasing number of loci, but little change in the decrease in diversity found at low migration rates (Fig. 4).

For genotypically redundant traits, we found that roughly 50% of the adaptive loci were highly differentiated between patches when migration was low (i.e. allele frequency differences of 95% or greater) (Fig. S9), and that there was little difference in the dd-slopes between divergent and uniform selection regimes across low migration high redundancy parameter space (Fig. S10).

Finally, mutation rate had little effect on any of the qualitative patterns that were seen over very low migration rates (Fig. S8). In contrast, across higher migration rates, increasing the mutation rate attenuated both the increase in diversity as well as the increase in divergence observed near the selected loci in the parameter sets with a small number of loci (Fig. S8).

### Single-Locus, Cline Model

We explored how more realistic models of population structure influenced patterns in nucleotide diversity and F_ST_ by investigating the previously-described metrics across a linear, ten-patch cline. We found that patches on the interior of the cline (i.e. patches 4-7) produced qualitative patterns very similar in nature to the two-patch model, including positive dd-slopes over low migration rates and negative dd-slopes over intermediate-high migration rates (Fig. 5a-b). Moving towards the exterior of the cline (i.e. patches 1-3 & 8-10), the dd-slopes were negative over a much reduced region of the migration-selection parameter space explored (Fig. 5a-b). In the patches at either end of the cline, the dd-slope was positive for all migration rates save for the very extremes (Fig. 5a-b). Similar to the two-patch scenario, the slope of F_ST_ by distance reached a maximum magnitude at an intermediate-high migration rate, before increasing to approximately zero beyond the critical migration rate (Fig. 5c).

**Figure 5:**
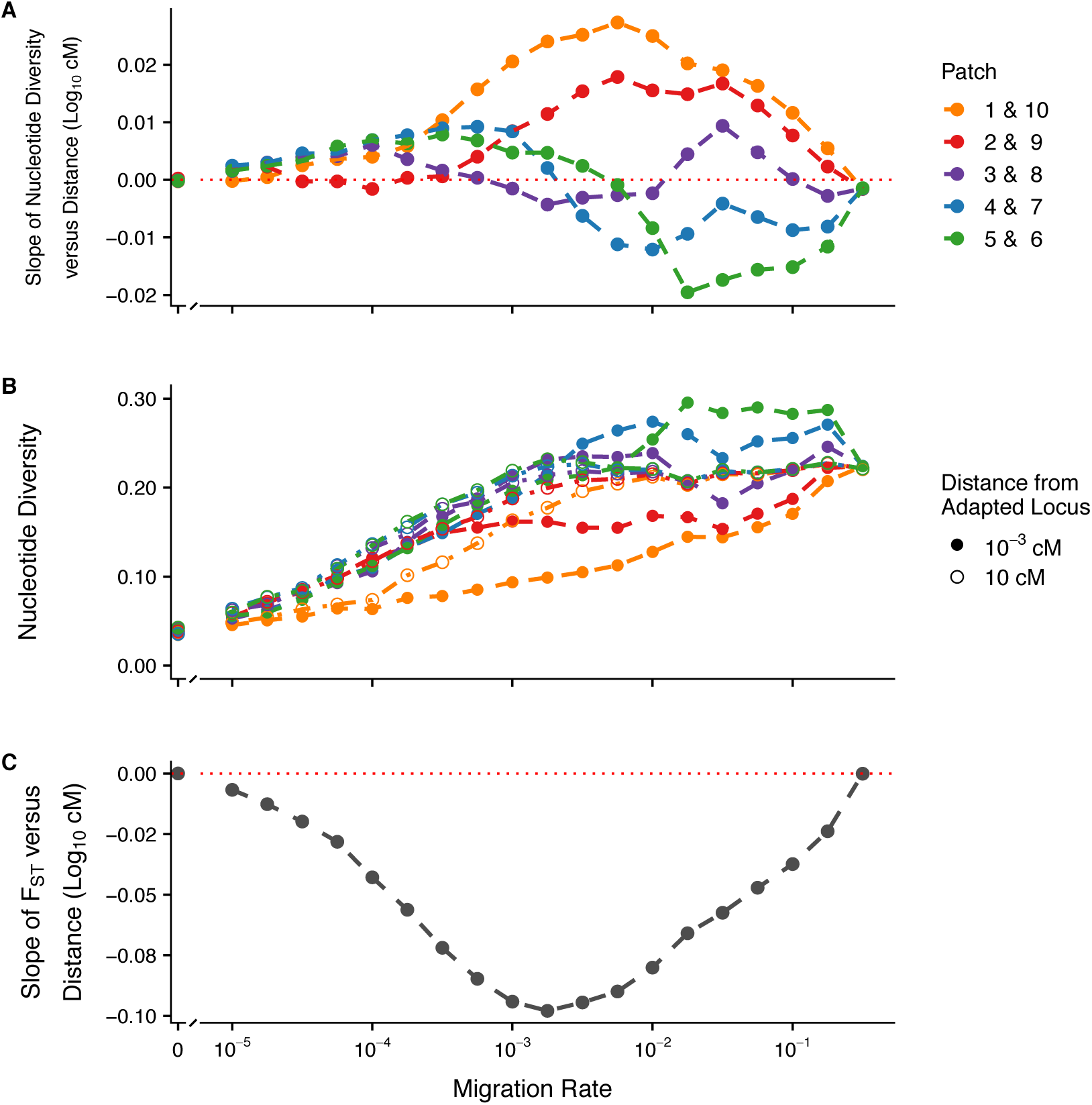
Effect of migration-selection balance on diversity at neutral sites linked to a single divergently selected locus in ten-patch cline model. Each patch was comprised of N = 1,000 individuals, mutation rate = 10^−5^ per locus, the V_S_ = 5. Panels are as described in Fig. 1.

## DISCUSSION

### Local adaptation can cause peaks or troughs in nucleotide diversity

To study how local adaptation in heterogenous environments shapes patterns in nucleotide diversity we assessed the diversity at neutral sites linked to a causal locus driving a trait under spatially divergent selection over a wide range of parameter space. Broadly, we demonstrate that no single signature for nucleotide diversity within populations is characteristic of local adaptation at equilibrium. As most previous work focused primarily on studying among-population diversity as the primary signature of local adaptation, this helps contextualize the contrasting results about nucleotide diversity found in the literature (Petry 1983; Nordborg 1997; Charlesworth et al. 1997; Sakamoto and Innan 2019). We now summarize how these evolutionary processes interact to yield these contrasting patterns of within-population diversity. When migration is sufficiently low that migrant haplotypes don’t persist for long, and are rapidly selected out of the population, local adaptation produces a pattern resembling background selection (Fig. 1a, positive region, as per Nordborg 1997). Put another way, when migration is very weak, nucleotide diversity for a very tightly linked neutral locus tends towards the expectation for a single patch without migration, as selection is so strong that the effective migration rate it experiences is near 0 (as per Barton and Bengtsson 1986). Thus, the effective population size it experiences tends towards that of a single population of the same size without migration. In the two-patch model here, this results in nucleotide diversity ∼1.5x lower than at the unlinked locus (Fig. 1b), which experiences an effective population size more congruent with that of the metapopulation as a whole (Whitlock and Barton 1997).

The widths of the regions of depleted diversity observed here are similar to the expectations for background selection (as per Nordborg 1997), where diversity is predicted to be depleted on the order of 10’s of cM away from the causal locus under similar conditions (Hudson and Kaplan 1995). By contrast, with selective sweeps, diversity is predicted to be depleted more deeply, but over a narrower region, on the order of centiMorgans away from a focal locus (Barton 2000). While the width of the region observed here is similar to the expected width under background selection (Fig. S3a), the magnitude of depletion is substantially greater with local adaptation (Fig. 1a). The effect of background selection can be seen when *m* = 0, as both populations evolve to their respective equilibria and further mutations are deleterious and are selected against. Comparing the case with *m* = 0 to cases with low migration, diversity is much more strongly reduced in the latter (Fig. 1a), as the rate that negatively selected alleles are introduced into the population is greater (*m* > μ) and because the magnitude of the effect size of those alleles is, on average, greater.

Based on these results, we would predict that particularly extreme reductions in nucleotide diversity at linked sites might be seen in small peripheral populations experiencing weak migration and strong selection. In this case, neutral regions of the genome would have levels of nucleotide diversity similar to those expected for the effective size of the metapopulation, whereas loci linked to the selected locus would have levels similar to those expected for the effective size of the small peripheral population, which could be much more discordant than found with the symmetrical population sizes simulated here.

In the above described parameter space, where we find an erosion of within-population diversity at the neutral loci flanking locally adapted loci, we also find an increase in F_ST_ at the same sites (Fig 1c). Thus, this pattern which has been interpreted as a result of background selection or uniform positive selection (Noor and Bennett 2009; Cruickshank and Hahn 2014), can also be driven by migration-selection balance, as could be predicted from older migration-selection studies (e.g. Petry 1983; Bengtsson 1985; Bengtsson and Barton 1986). Our results do not discount the effects positive selection or purifying selection may have on producing signatures resembling the classic “genomic islands of differentiation” (e.g. Via 2009; Nosil 2009), but do show that local adaptation could also generate similar patterns in nucleotide diversity and F_ST_.

Conversely, when migration is higher but not so strong as to collapse the locally adapted polymorphism, we find peaks in both within-population nucleotide diversity (Fig. 1a, negative region) and F_ST_ (Fig. 1c) at the neutral loci flanking locally adapted loci, although this effect is attenuated with larger population size (Fig. 2). Selection generates linkage disequilibrium between neutral loci and the locally adapted locus proportional to the recombination distance between them; when the locally adapted haplotype migrates into its maladapted patch the diversity is transiently increased at the flanking neutral loci. When locally adapted haplotypes migrate into the maladapted patch at a greater rate than selection can effectively purge them, sharp peaks are generated in both within-population diversity and F_ST_. These peaks attenuate with larger population sizes, as the maintenance of linkage disequilibrium is increased over a greater range of recombination with smaller populations (as per Ohta and Kimura 1971). While a limited effect of increased diversity was noted by Sakamoto and Innan (2019), this was not discussed as a potentially important signature of local adaptation. Here, we show that considerable increases in nucleotide diversity can be found, especially when selection is strong, migration rate high, and effective population sizes are small, as is the case in many empirical examples of local adaptation. Concurrent peaks in within-population diversity and F_ST_ may therefore constitute an important signature of local adaptation that can be readily distinguished from background and positive selection – not all local adaptation will cause increased diversity along with peaks in F_ST_, but the detection of such patterns strongly suggests local adaptation as a driving process.

It is perhaps interesting to contrast these results with the expectation for standing genetic variation (V_G_) in a quantitative trait, which shows an increase in V_G_ under high migration rates, but no reduction in V_G_ under low migration rates (McDonald and Yeaman 2018). This further highlights the importance of clearly specifying model expectations, and the problems inherent in using nucleotide diversity data as a proxy for V_G_ (as per Reed and Franhkam 2001). Clearly, the question of “what maintains variation?” has a very different meaning for different kinds of variation.

### Effects of genotypic redundancy

For traits that have no genotypic redundancy, the effect of selection on the phenotype is divided among loci in proportion to their effect sizes (Yeaman 2015). As such, the strength of selection per locus is reduced as the number of loci increases, and the peaks and troughs in diversity at linked sites also attenuate, as described above (Fig 3). In contrast, when traits are genotypically redundant, we still find strong trough-like patterns in linked diversity as the number of loci increases (Fig. 4). When migration is low (*m* << 1/N_e_), different populations evolve essentially as though they were independent of one another, with each population converging on a certain combination of alleles that achieves its phenotypic optimum effectively independent of the particular combination of alleles the other population is using. As a result, the two different populations tend to evolve very different combinations of alleles and end up differentiated at substantially more loci than would strictly be needed to achieve local adaptation in each patch (Fig. S9). When the different populations are differentiated at such a large proportion of loci, matings between divergently adapted individuals result in F2 hybrid breakdown (Yeaman and Whitlock 2011; Thompson et al. 2019), causing selection to operate on many more loci than would be expected solely due to local adaptation. The action of selection on all differentiated loci thereby causes the background selection-like effect across many loci, and prevents the attenuation of the trough-like pattern at low migration rates and many redundant loci. Furthermore, the population structure that is generated through low migration and high genotypic redundancy (as per Goldstein and Holsinger 1992; Phillips 1996) generates a similar trough pattern even with spatially uniform selection (Fig. S10). If many genotypically redundant loci scattered throughout the genome contribute to local adaptation, this could cause an extensive decrease in nucleotide diversity at low migration rates. Studying the prevalence of F2 hybrid breakdown among divergent populations or sub-species could help indicate whether highly redundant traits causing this kind of effect are common.

### Peaks and troughs in diversity under more realistic models of population structure

When a species range spans an environmental gradient, populations that inhabit the interior regions experience very different evolutionary processes from those inhabiting the periphery, due to variations in migration rate combined with the different combinations of allele frequencies found in different regions. We can see the effects of variable evolutionary processes over the species range in our cline model, where patterns in linked nucleotide diversity are starkly different between the populations that inhabit the interior of the range and those that inhabit the periphery (Fig. 5). Populations that inhabit the interior of the range have patterns in diversity that are qualitatively quite similar to those described in our two-patch model; in contrast, populations that inhabit the periphery present with depleted diversity across a substantially larger region of migration-selection parameter space and the sharp peaks in diversity seen in our two-patch results do not appear (Fig. 5).

Across a spatially extended clinal environment, the interior populations receive an influx of maladapted alleles at a relatively high rate (here, ≳2x compared to the peripheral populations), which increases the nucleotide diversity at flanking loci in these populations. While migration is introducing differently adapted alleles into the interior populations, however, selection is purging them. The purging of maladapted alleles across the interior populations, coupled with the stepping stone nature of our model, ultimately results in the decreased influx of maladapted alleles into the peripheral populations (i.e. few haplotypes with an allele optimally adapted to one end of the cline migrate to the other end). As such, the degree of polymorphism that is maintained around the locally adapted locus in the peripheral populations is not sufficient enough to produce the peaks in diversity seen in our two-patch results. Consequently, when species adapt over spatially extended clinal environments, it may be very possible to find both peaks and troughs in nucleotide diversity at a single locus within a single species.

## CONCLUSION

We demonstrate that there is no universal nucleotide-scale signature of local adaptation, even with the simplest possible model of spatially divergent selection. Nucleotide diversity within populations can be substantially decreased or increased depending on the relative strengths of migration and selection. Additionally, local adaptation can result in regions of depleted within-population diversity over chromosomal distances similar to that of background selection, with a substantially greater magnitude of diversity eroded than with background selection. Our results demonstrate that local adaptation must also be considered, in addition to background selection and selective sweeps, when making inferences based on genomic regions of reduced diversity. While reductions in diversity are not particularly diagnostic, peaks in nucleotide diversity are only expected under local adaptation or other models of balancing selection, and as such, can distinguish local adaptation vs. uniform positive/purifying selection (as per Booker et al. 2019). Finally, our results from models with increased realism further highlight that there is little reason to expect a consistent pattern in nucleotide diversity across heterogeneous environments, as patterns of decreased or increased diversity can be expected depending on polygenicity, redundancy, geography, migration, and selection.

## AKNOWLEDGEMENTS

We would like to thank S. Aeschbacher and M. Nordborg for constructive discussion of the literature, M. Williamson and M. Whitlock for assistance reconstructing previous analytical solutions, and T. Booker, C. Rougeux, and M. Whitlock for their constructive feedback on the manuscript. This project was enabled in part by computational support provided by Compute Canada, and was funded by an NSERC scholarship to RJ, an NSERC Discovery grant to SY, and an AIHS chair to SY.

