## Supplemental Information for "Local adaptation can cause both peaks and troughs in nucleotide diversity within populations"

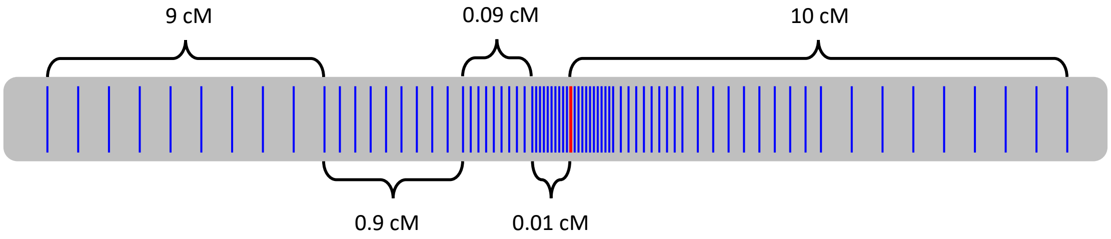


**Figure S1: Genetic map of chromosome with a single divergently selected locus.** The chromosome consisted of one divergently selected locus and 74 neutral loci. The selected locus is denoted by the red bar, the neutral loci are denoted by the blue bars. The neutral loci were positioned about the selected locus symmetrically at distances from $10^{-3}$ to $10^{1}$ cM away on a $\log_{10}$ scale.

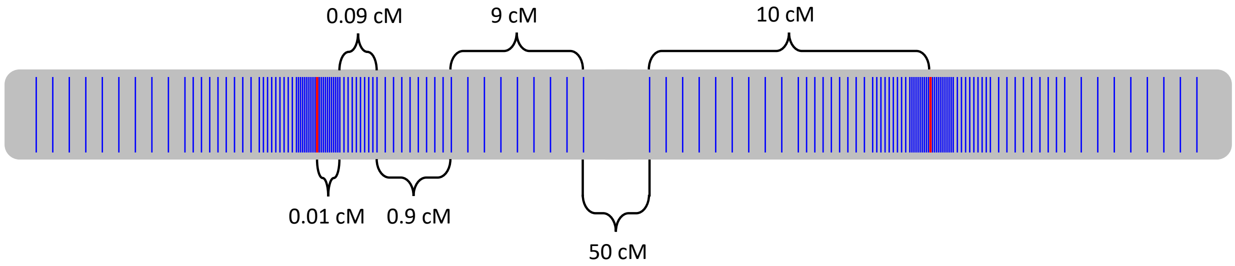


**Figure S2: Genetic map of chromosome with multiple divergently selected loci.** Each complement of a selected locus and its 74 flanking neutral loci was separated by at least 50 cM. All genetic architectures with more than a single selected locus followed this same pattern.

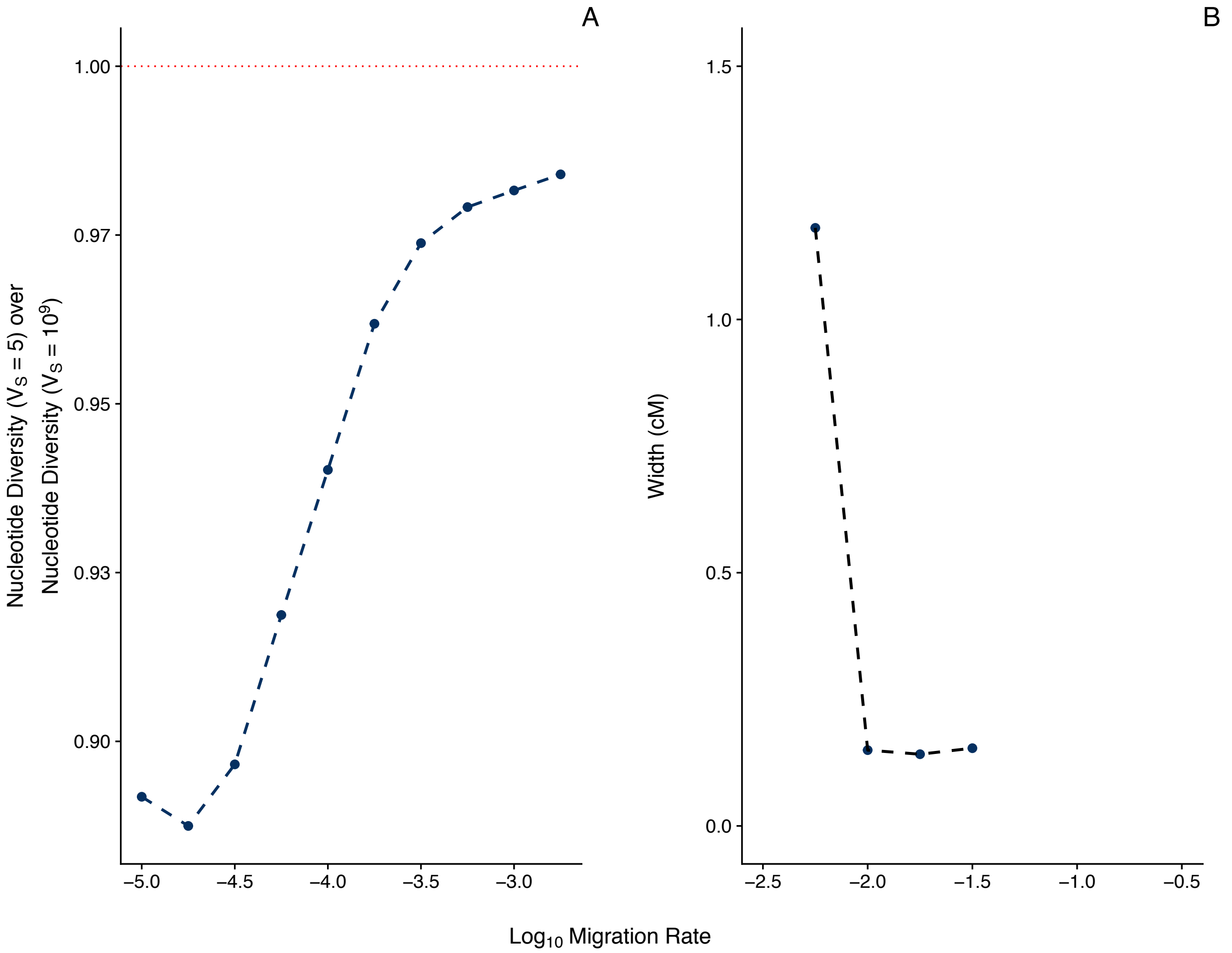


**Figure S3: Genetic distance over which diversity at neutral sites is affected by linkage to a single divergently selected locus in two-patch model.** Panel A shows the degree to which within-population nucleotide diversity is eroded at the neutral loci 9 to 10 cM away from the locally adapted locus under strong selection ($V_S = 5$) relative to genome-wide background levels ($V_S = 10^9$). Panel B shows the width (cM) of peaks in within-population nucleotide diversity at the neutral loci linked to the locally adapted locus. Each patch was comprised of N = 1,000 individuals and the mutation rate = $10^{-5}$ per locus.

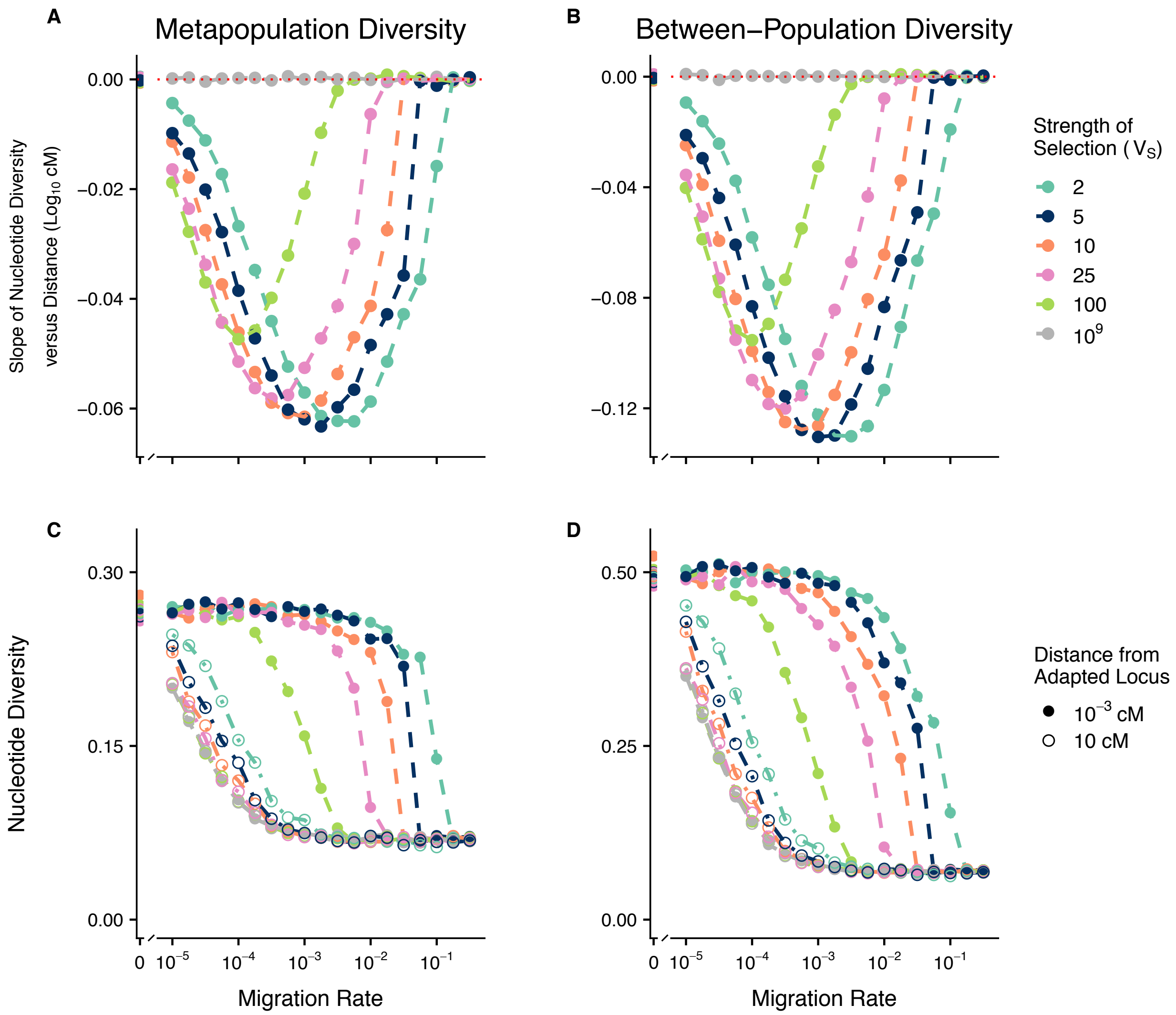


**Figure S4: Effect of migration-selection balance on total metapopulation diversity and between-population diversity ($d_{xy}$) at neutral sites linked to a single divergently selected locus in two-patch model.** The slopes of metapopulation nucleotide diversity versus distance ($\log_{10}$ cM) (A) and between-population nucleotide diversity versus distance ($\log_{10}$ cM) (B), and the metapopulation nucleotide diversity (C) and between-population nucleotide diversity (D) at the loci closest ($10^{-3}$ cM) and furthest (10 cM) from the locally adapted locus are shown against the $\log_{10}$ migration rate after 50,000 generations. Each patch was comprised of N = 1,000 individuals and the per-locus mutation rate = $10^{-5}$.

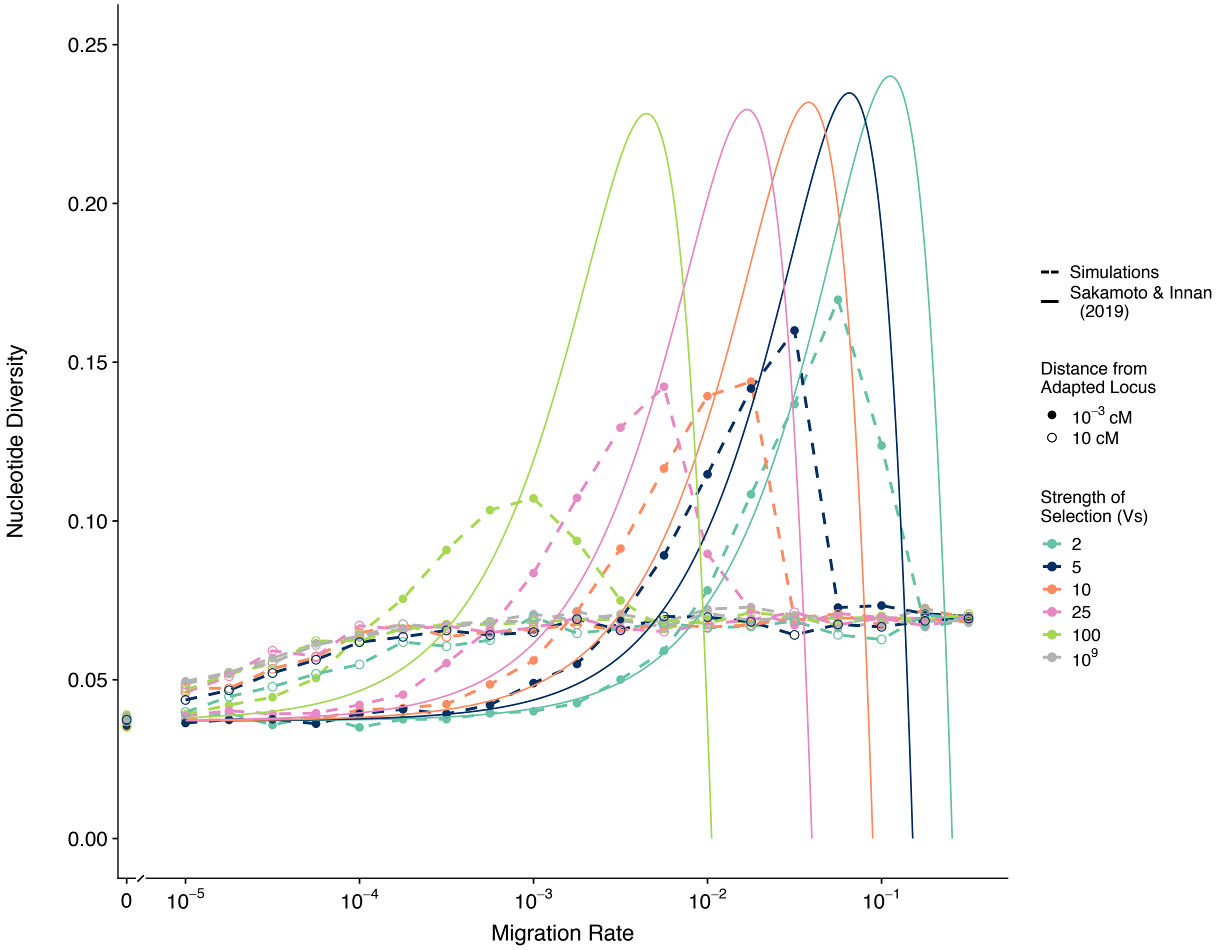


**Figure S5: Comparison between simulations results and analytical predictions at a neutral site linked to a single divergently selected locus in two-patch model.** Simulation results for the within-population nucleotide diversity at a neutral locus $10^{-3}$ cM from a divergently selected locus are shown against the $\log_{10}$ migration rate after 50,000 generations. Each patch was comprised of N = 1,000 individuals and the per-locus mutation rate = $10^{-5}$. Analytical predictions for the expected within-population heterozygosity at a similarly linked neutral locus from Sakamoto and Innan (2019) equation 25 are shown for comparison. Analytical predictions were generated using symmetrical selection coefficients to parameterize eq. 25 (i.e. their $s_1 = |s_2|$). Simulation results are shown in dashed lines, analytical predictions are shown in solid lines.

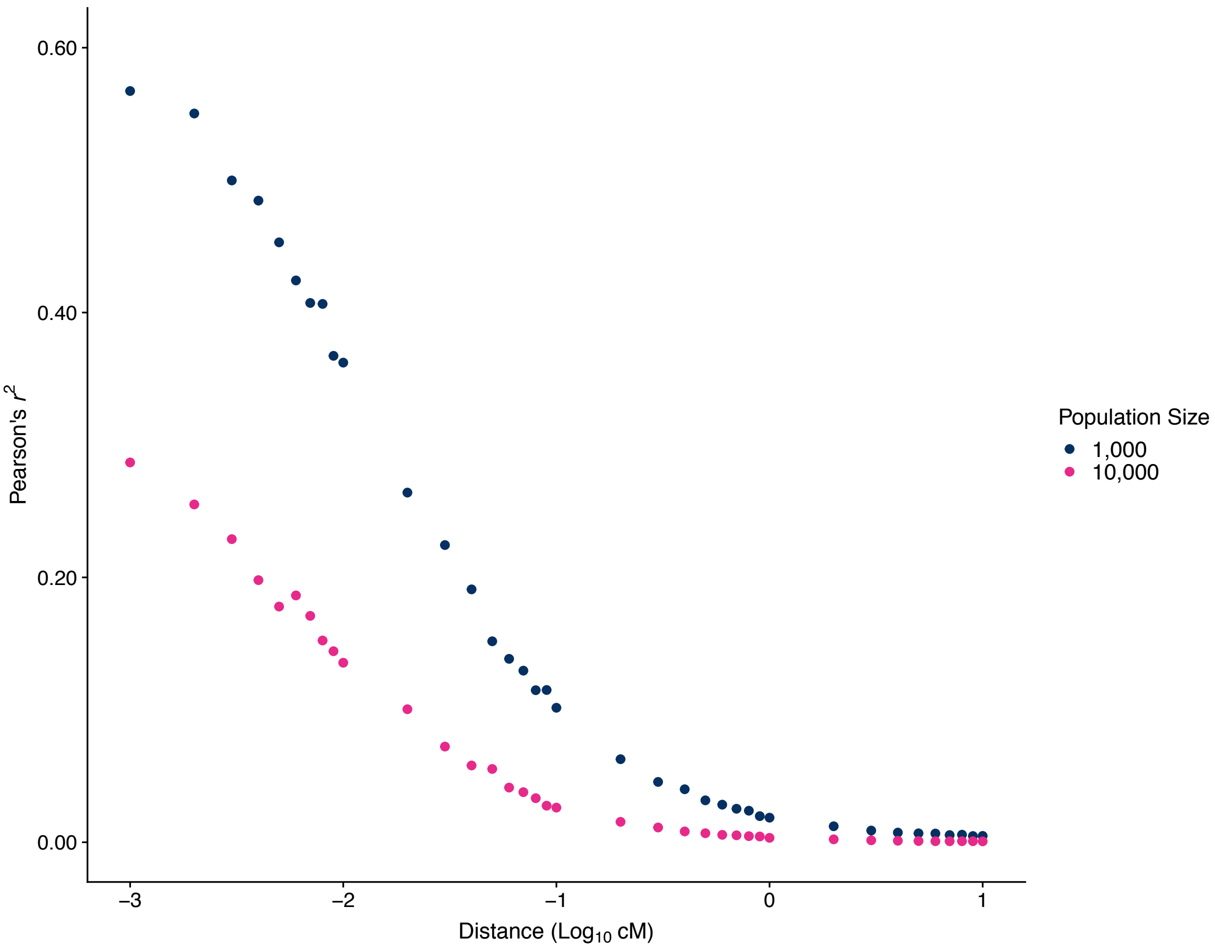


**Figure S6: Effect of population size on linkage disequilibrium between neutral loci and a single divergently selected locus in two-patch model.** A migration rate of $10^{-1.5}$, mutation rate of $10^{-5}$, and $V_S$ of 5 is shown.

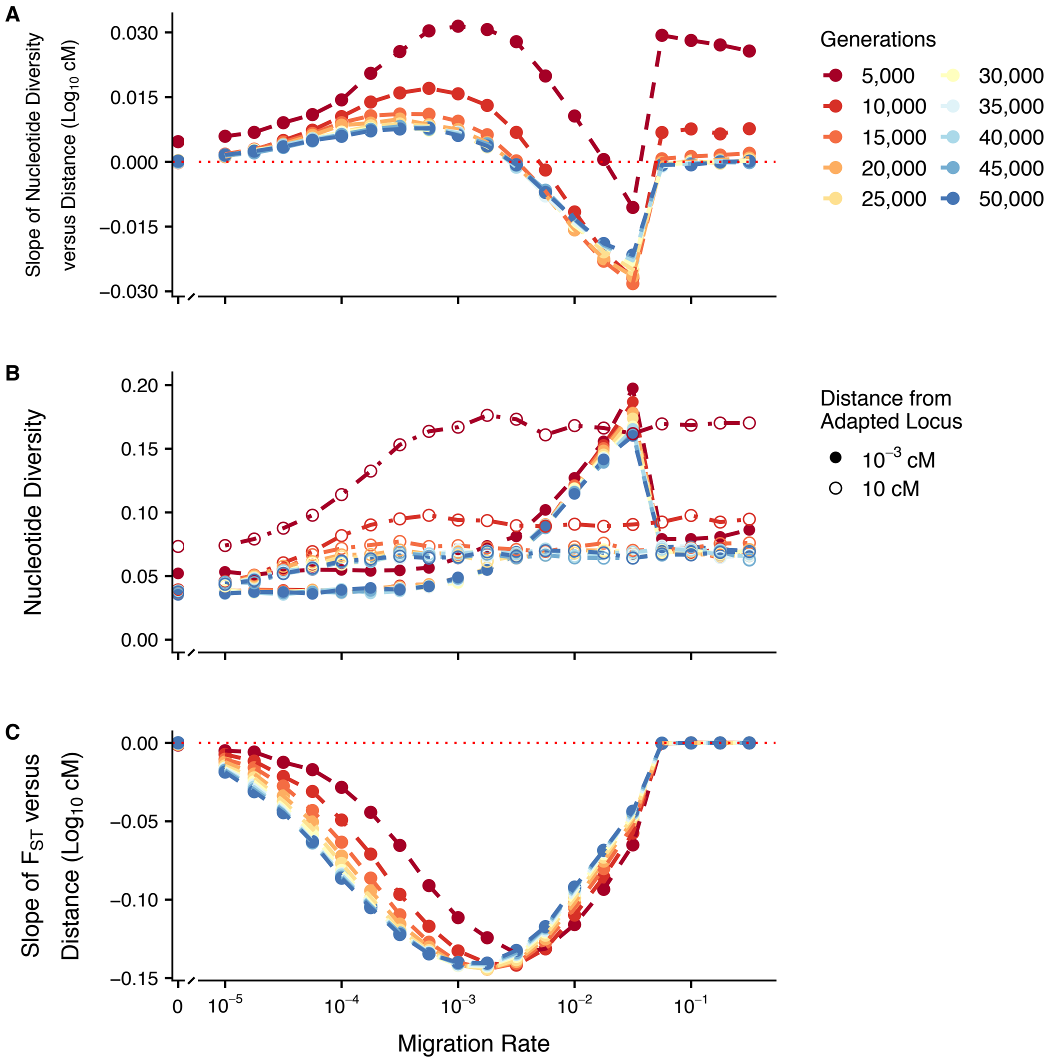


**Figure S7: Time series effect of migration-selection balance on diversity at neutral sites linked to a single divergently selected locus in two-patch model.** Each patch was comprised of N = 1,000 individuals, mutation rate per locus = $10^{-5}$, and $V_S = 5$. Panels are as described in Fig. 1 in 5,000 generation intervals.

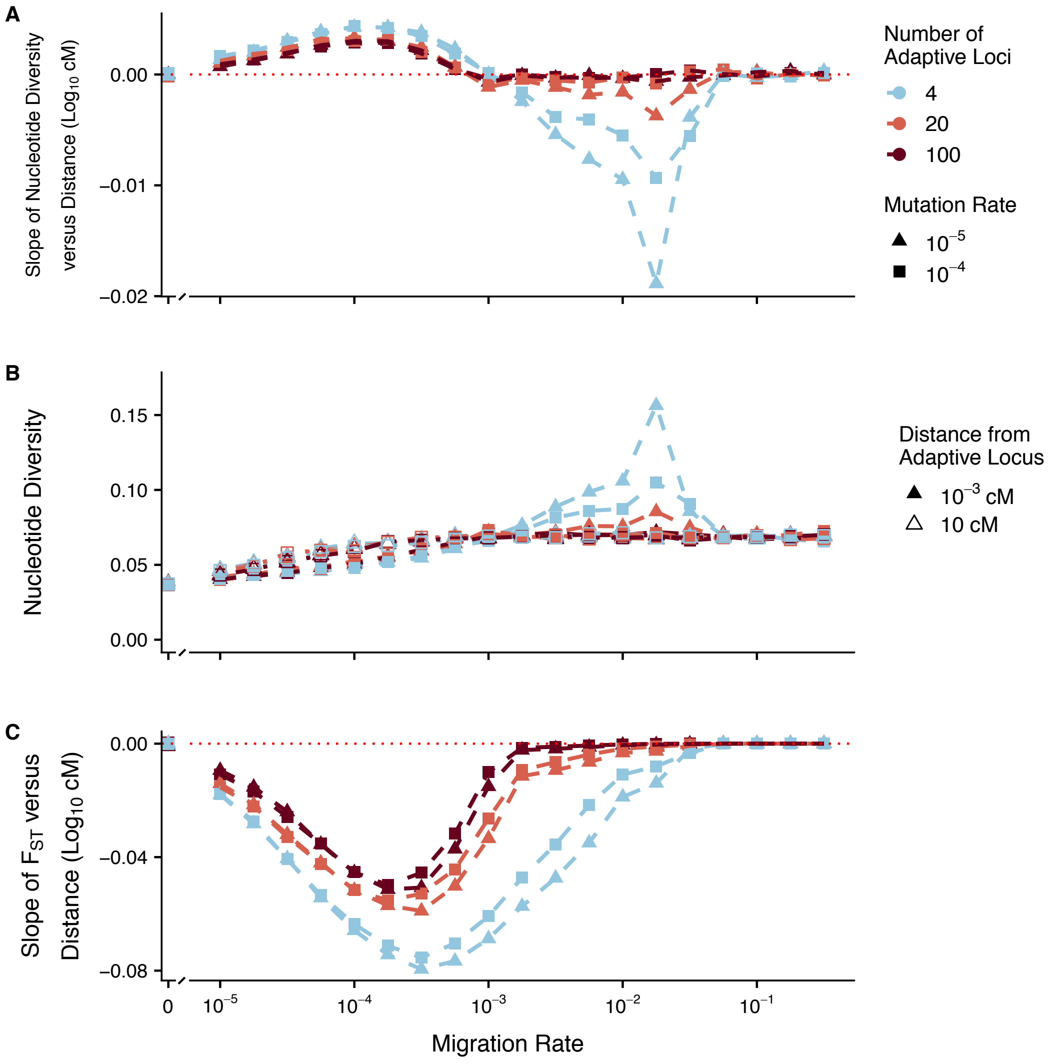


**Figure S8: Effect of migration-selection balance and adaptive mutation rate on diversity at linked neutral sites for a quantitative trait with different numbers of loci and variable levels of genotypic redundancy.** Allele effect sizes were ± 0.25, such that an individual could reach the optimum in a patch (± 1) by being homozygous for the optimal allele at 2 loci. Mutation rate at selected loci = $10^{-5}$ or $10^{-4}$, and at neutral loci = $10^{-5}$. Each patch was comprised of N = 1,000 individuals and $V_S$ = 5. Panels are as described in Fig. 1.

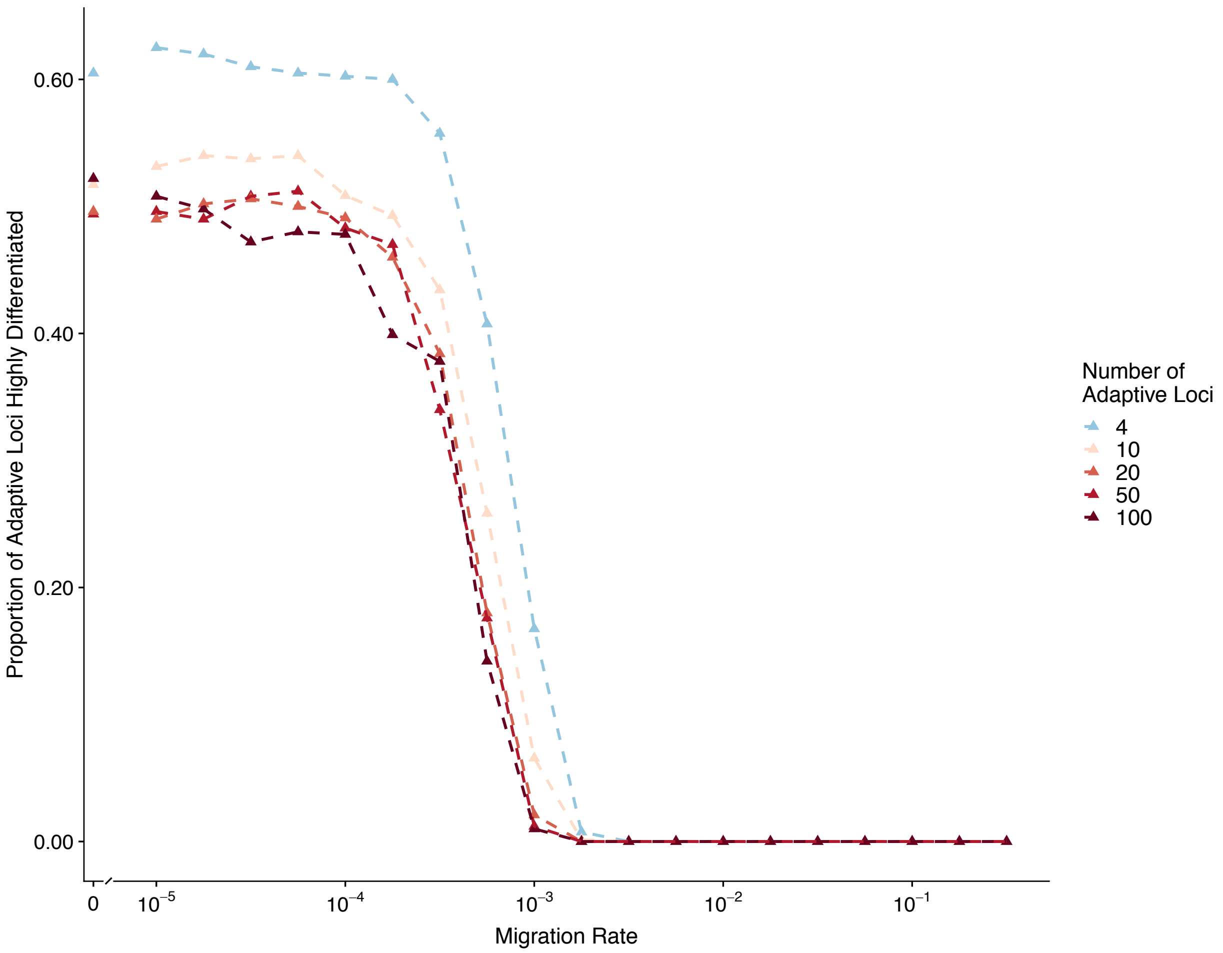


**S9 Proportion of adaptive loci highly differentiated between patches for a quantitative trait with different numbers of loci and variable levels of genotypic redundancy.** Loci with allele frequency differences of 95% or greater between patches were deemed highly differentiated. Allele effect sizes were ± 0.25, such that an individual could reach the optimum in a patch (± 1) by being homozygous for the optimal allele at 2 loci. Results after 50,000 generations shown; each patch was comprised of N = 1,000 individuals, mutation rate = $10^{-5}$ per locus, and $V_S = 5$.

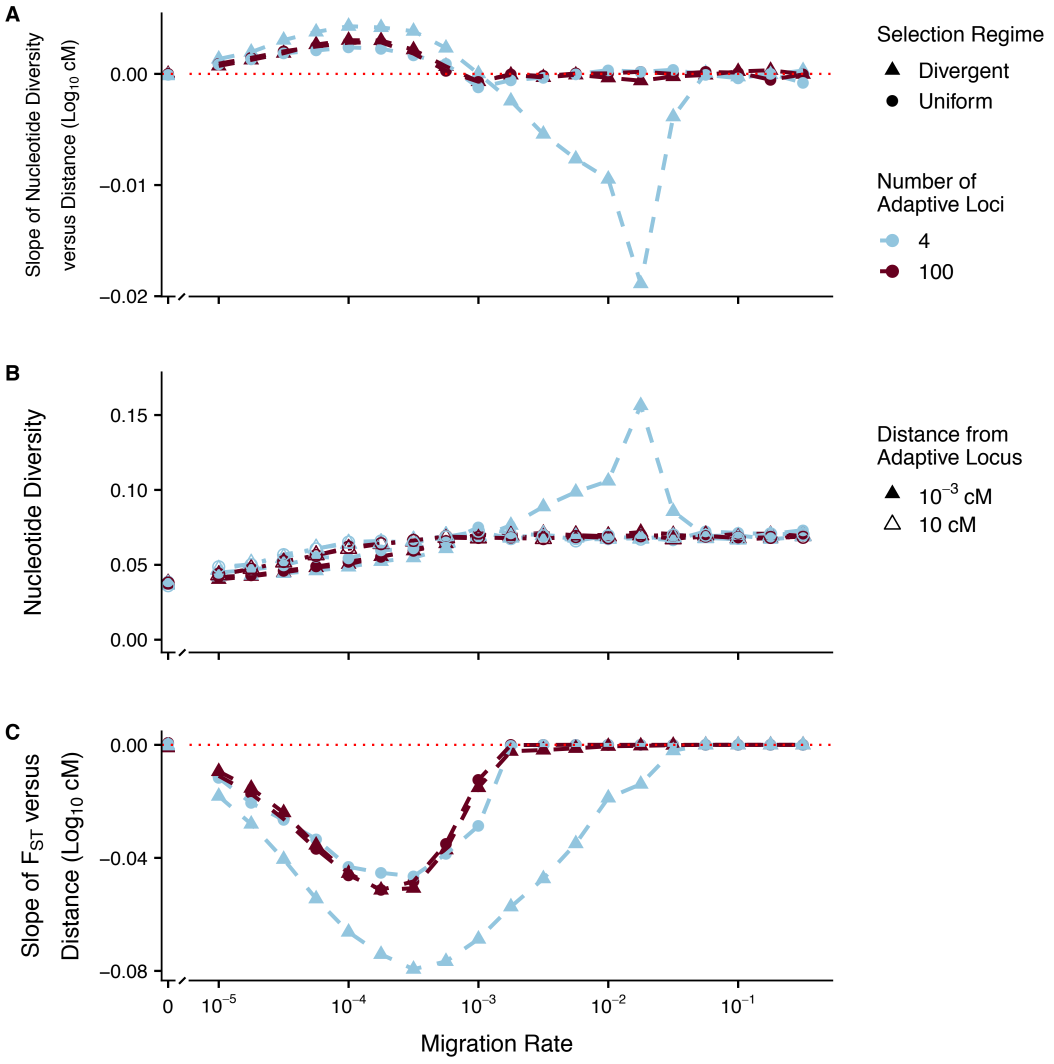


**S10 Effect of migration-selection balance and selection regime on diversity at linked neutral sites for a quantitative trait with different numbers of loci and variable levels of genotypic redundancy.** Allele effect sizes were ± 0.25, such that an individual could reach the optimum in a patch (± 1 divergent selection; +1 uniform selection) by being homozygous for the optimal allele at 2 loci. Panels are as described in Fig. 1; each patch was comprised of N = 1,000 individuals, mutation rate = $10^{-5}$ per locus, and $V_S$ = 5.
